# Calmodulin acts as a state-dependent switch to control a cardiac potassium channel opening

**DOI:** 10.1101/2020.07.04.187161

**Authors:** Po Wei Kang, Annie M. Westerlund, Jingyi Shi, Kelli McFarland White, Alex K. Dou, Amy H. Cui, Jonathan R. Silva, Lucie Delemotte, Jianmin Cui

## Abstract

Calmodulin (CaM) and PIP_2_ are potent regulators of the voltage-gated potassium channel KCNQ1 (K_V_7.1), which conducts the I_Ks_ current important for repolarization of cardiac action potentials. Although cryo-EM structures revealed intricate interactions between the KCNQ1 voltage-sensing domain (VSD), CaM, and PIP_2_, the functional consequences of these interactions remain unknown. Here, we show that CaM-VSD interactions act as a state-dependent switch to control KCNQ1 pore opening. Combined electrophysiology and molecular dynamics network analysis suggest that VSD transition into the fully-activated state allows PIP_2_ to compete with CaM for binding to VSD, leading to the conformational change that alters the VSD-pore coupling. We identify a motif in the KCNQ1 cytosolic domain which works downstream of CaM-VSD interactions to facilitate the conformational change. Our findings suggest a gating mechanism that integrates PIP_2_ and CaM in KCNQ1 voltage-dependent activation, yielding insights into how KCNQ1 gains the phenotypes critical for its function in the heart.

## Introduction

The KCNQ (K_V_7) voltage-gated potassium channel family consists of five members (KCNQ1-KCNQ5) which provide K^+^ currents critical for physiological function of both excitable and non-excitable tissues (1–5). All KCNQ channels require the association of calmodulin (6, 7), and membrane lipid PIP_2_ (8, 9) for voltage dependent opening. In the heart, the predominant cardiac isoform KCNQ1 (K_V_7.1) also associates with the auxiliary subunit KCNE1 (1, 2) to carry the slow delayed rectifier K^+^ current (I_Ks_). I_Ks_ facilitates repolarization of the cardiac action potential, especially during β-adrenergic stimulation (10, 11). Hundreds of mutations in KCNQ1 are linked to arrhythmias such as atrial fibrillation and long QT syndrome (LQTS), predisposing patients to sudden cardiac death (4, 12, 13).

Structurally, KCNQ1 channels comprise four identical subunits, each with six transmembrane (S1-S6) and four cytoplasmic (helix A to D; HA-HD) α-helices, with S1-S4 forming the voltage-sensing domain (VSD) and S5-S6 the pore domain (14, 15) (Fig. 1A-B). Upon membrane depolarization, the KCNQ1 VSDs activate with two experimentally resolvable steps: first from the resting state to an intermediate state, and then to the fully-activated state (8, 16–22). The VSD conformational changes are communicated to the pore domain via a series of residue-residue interactions, leading to pore opening (16, 18). KCNQ1 features a unique two-open-state gating mechanism (Fig. 1C). Previous work showed that KCNQ1 VSD can trigger pore opening when activated to both the intermediate and the fully-activated states, yielding two open states: the intermediate-open (IO) and activated-open (AO) states (8, 18, 21, 23, 24). The IO and AO states have been shown to give rise to bi-exponential current activation kinetics (18, 23, 24), with the fast and slow components approximating the IO and AO states, respectively (Fig. 1C). The IO and AO states exhibit different functional properties important for KCNQ1 physiological function, with regulation of IO and AO by different auxiliary subunits leading to tissue specific current phenotypes (21). For example, although KCNQ1 conducts predominantly at the IO-state as seen by the large fraction of the fast-current component (Fig. 1C), the auxiliary subunit KCNE1 eliminates the kinetically fast IO state and enhances the slow AO state, contributing to the kinetically slow I_Ks_ suitable for cardiac tissue needs (8, 18, 21).

**Figure 1.**
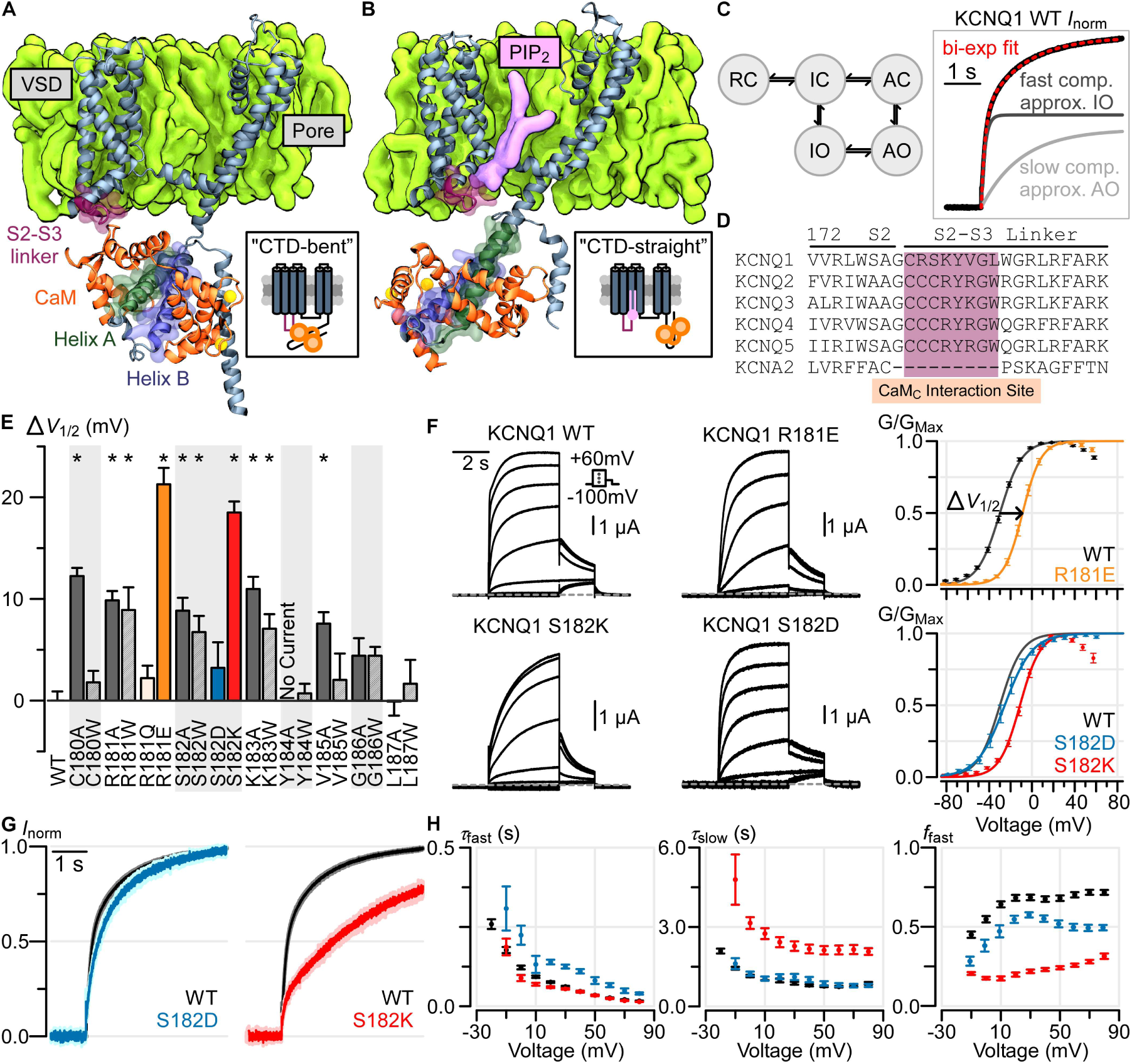
The S2-S3 linker within the KCNQ1 voltage-sensing domain modulates KCNQ1 currents. **(A-B)** Structural depiction of one monomer of the tetrameric KCNQ1-CaM complex when solved in the absence (A) and presence (B) of PIP_2_ (ie. “CTD-bent” (A) and “CTD-straight” (B) conformations). **(C)** Left: A simplified KCNQ1 gating model (8, 18, 21, 23, 24) with two open states, intermediate-open (IO) and activated-open (AO). Horizontal transitions correspond to VSD activation, vertical transitions correspond to pore opening. Abbreviations: RC: Resting-Closed, IC: Intermediate-Closed, AC: Activated-Closed, IO: Intermediate-Open, AO: Activated-Open. Right: Exemplar bi-exponential activation kinetics fitting with fast and slow components of a Q1-WT normalized current: voltage stepped from −80 mV to 0 mV. **(D)** Sequence alignment of the KCNQ family and KCNA2 at the S2-S3 linker region predicted to interact with CaM. **(E)** Mutagenesis screen of the KCNQ1 S2-S3 linker region shown in panel D. Y-axis: Δ*V*_1/2_ from WT channel (see panel F for example). X-axis: residue position. Stars indicate significant Δ*V*_1/2_ from WT (Supplementary table 1). Error bars are SEM. **(F)** Left: exemplar recordings from KCNQ1 WT and S2-S3 linker mutant channels. Voltage protocol is shown in inset. Currents were collected with 10-mV increments at the test pulse but shown in 20-mV increments to avoid crowding. Right: Average G-V relationship calculated for the respective channels. An example of Δ*V*_1/2_ measurement is shown for mutant R181E. **(G)** Average normalized KCNQ1 channel activation kinetics, stepping from −80 mV to 0 mV, for Q1-WT (black, *n* = 10), Q1-S182D (blue, *n* = 7), and Q1-S182K (red, *n* = 10) channels. The light shadings show SEM. (F) Average bi-exponential kinetic fit parameters for Q1-WT (black), Q1-S182D (blue), and Q1-S182K (red) channels. All current traces at each test voltage were first fitted with a bi-exponential equation and normalized by the fitted steady-state current amplitude (panel G). Error bars are SEM. τ_*fast*_ and τ_slow_ are the time constants for the fast and slow components, respectively. f_fast_ is fraction of total current contributed by the fast component.

KCNQ regulation by calmodulin (CaM) (6, 7, 14, 15) is still not clearly understood. CaM features a two-lobe structural architecture with the N-terminal domain (N-lobe) and the C-terminal domain (C-lobe), and each lobe harbors two Ca^2+^ binding sites (25). CaM regulation of KCNQ channels is still debated, some studies suggested that CaM confers Ca^2+^-dependent inhibition of KCNQ2/3 and KCNQ4 currents but Ca^2+^-dependent enhancement of KCNQ1 currents (26–32). Other studies found that CaM C-lobe bound to KCNQ1 does not coordinate Ca^2+^ ions under 1 mM EGTA or 5 mM Ca^2+^ conditions (33). CaM binding by itself may therefore be sufficient to affect KCNQ1 function, similar to other voltage-gated channels (34, 35). Structurally, CaM binds the KCNQ1 C-terminus domain (CTD) such that CaM N-lobe associates with KCNQ1 Helix B (HB) and CaM C-lobe interacts with KCNQ1 Helix A (HA) (Fig. 1A) (6, 7, 14, 15, 33). Recent cryo-EM structural studies of the full-length KCNQ1/CaM complex (14, 15) revealed intricate interactions between CaM, the KCNQ1 VSD, and the membrane lipid PIP_2_. In the absence of PIP_2_, the KCNQ1 CTD “bends” at the S6-HA linker to enable CaM interaction with the VSD S2-S3 linker (Fig. 1A) (14, 15). In the presence of PIP_2_, the KCNQ1 CTD “straightens” to disengage CaM from the VSD, and the S2-S3 linker instead interacts with PIP_2_ (Fig. 1B) (15). We will refer to these two cryo-EM structural states as the “CTD-bent” and the “CTD-straight” conformations (Fig. 1A-B). The VSDs in both conformations adopt the fully-activated state (14, 15, 21). The CTD-bent conformation features a closed pore, while the CTD-straight conformation exhibits a dilated pore (14, 15).

Although the KCNQ1/CaM structures provided new insights to the conformational ensemble of KCNQ1, it is still unknown how these structures correlate to the channel’s voltage-dependent activation process. First, the CTD-straight structure likely corresponds to the AO state (Fig. 1C) based on its activated VSD and dilated pore (14, 15, 21), but functional data is lacking to support this. Moreover, the CTD-straight structure additionally features binding by the auxiliary subunit KCNE3 (15), which may lead to conformational differences compared to KCNQ1/CaM alone. Second, it is unknown whether the CTD-bent conformation corresponds to a functional state in the KCNQ1 activation pathway, since PIP_2_ exists under physiological conditions and the structure was solved without PIP_2_. By extension, it is unclear whether the CaM-VSD interactions seen in the CTD-bent conformation occurs during gating. In fact, no studies have shown simultaneous CaM and PIP_2_ binding to the KCNQ1 VSD. The available studies thus suggest mutually exclusive CaM or PIP_2_ binding to the S2-S3 linker (14, 15) (Fig. 1A-B), with PIP_2_ outcompeting CaM for the activated VSD. Third, if the CTD-bent structure represents a functional state in the KCNQ1 activation pathway, what is the role of CaM-VSD interactions in KCNQ1 gating? In summary, the relationship between KCNQ1/CaM structures and voltage-dependent gating, specifically the role of CaM-VSD interactions, remains unresolved.

Here, we combine electrophysiology experiments and network analysis of molecular dynamics (MD) simulations to show that CaM interacts with the KCNQ1 VSD during voltage-dependent activation to modulate ionic current, despite PIP_2_ presence in the membrane. We demonstrate that both the CTD-straight and CTD-bent conformations are relevant in the KCNQ1 voltage-dependent gating process. We further provide evidence that the transition between the CTD-bent and CTD-straight conformations during gating is driven by VSD state-dependent interactions between the S2-S3 linker, CaM, and PIP_2_. Functionally, we find that the loss of CaM-VSD interactions during gating control channel transition into the AO state, while having no effect on the IO state. Lastly, we map a CTD motif, formed by the S6-HA and HB-HC linkers, that works together with the CaM-VSD interface to affect pore opening to the AO state. Based on our results, together with the cryo-EM structural data, we suggest a gating model in which CaM act as a VSD state-dependent switch to alter pore opening via the KCNQ1 CTD, thereby tuning the slow kinetics related to the cardiac repolarizing currents.

## Results

The cryo-EM studies showed two KCNQ1 structures with large conformational differences, with the CTD-straight structure solved in the more physiological conditions featuring lipid nanodiscs in the presence of PIP_2_ and the CTD-bent structure solved in the absence of PIP_2_ (14, 15). The CTD-straight structure is likely to reflect the AO state based on its activated VSD and dilated pore (15), but whether the CTD-bent structure corresponds to any gating state in voltage dependent activation is not clear. We designed experiments and molecular dynamic simulations to address the following questions: 1) Is the CTD-bent structure part of the voltage dependent activation process? 2) Are there any functional evidence to support that the CTD-straight structure corresponds to the AO state? 3) What is the mechanism and role of the conformational switch between the CTD-bent and CTD-straight in the gating process of KCNQ1?

### The S2-S3 linker within the KCNQ1 voltage-sensing domain modulates KCNQ1 currents

We first examined the functional role of the S2-S3 linker within the KCNQ1 VSD, as the S2-S3 linker interacts with either CaM or PIP_2_ in the cryo-EM structures (14, 15). To this end, we undertook a dual alanine/tryptophan mutational scan in the region of the S2-S3 linker predicted to interact with CaM or PIP_2_ (Fig. 1A-B, maroon). This region of the S2-S3 linker is conserved among the KCNQ family (Fig. 1D), but not in the wider K_V_ superfamily. We assayed the mutant channels by two-electrode voltage clamp (TEVC) in *Xenopus* oocytes, in which the oocytes endogenously supplied CaM and PIP_2_. Each mutant channel was characterized by their half-activation voltage (*V*_1/2_) of the measured conductance-voltage (G-V) relationship, which indicates the voltage dependence of channel open probability. The mutational scan revealed significant depolarizing shifts in the *V*_1/2_ when mutating residues C180-K183 and V185, but not residues Y184 and G186-L187 (Fig. 1E, Supplementary Table 1). These results show the functional impact of S2-S3 linker residues C180-K183 on KCNQ1 activation. We further found charge-dependent effects on KCNQ1 currents at positions R181 and S182: the mutants R181E and S182K induced ∼20 mV depolarizing shifts in the *V*_1/2_ (Fig. 1F, yellow and red). Conversely, the mutants R181Q and S182D caused little change from the wild-type (WT) G-V relationship (Fig. 1E-F, blue). As previous functional studies and the CTD-straight structure suggested that residue R181 interacts with PIP_2_ (15, 36), the functional effects of the R181 mutants is likely due to altered PIP_2_ binding. On the other hand, the CTD-bent structure showed that S182 interacts with CaM (14, 15); therefore, the S182 mutation effects may be due to disrupted CaM-VSD interactions.

We also noted that the mutant S182K exhibited significant slowdown in the current activation kinetics compared to the WT channels (Fig. 1G-H). As the fast and slow components of the KCNQ1 current kinetics approximate the IO and AO states (Fig. 1C), we quantified current activation by fitting a bi-exponential function. The fitted time constants (τ_fast_ and τ_slow_), as well as the fraction of total current carried by the fast vs. the slow component (*f*_fast_), were used to quantify current activation kinetics. The S182D mutant exhibited similar kinetics as the WT channel with small changes in τ_fast_ and *f*_fast_ compared to WT (Fig. 1G-H), which correlated with the minimal shift in *V*_1/2_ induced by the S182D mutation. By contrast, the S182K mutant revealed a significant slowing in current activation kinetics compared to WT channels (Fig. 1G-H), which correlated with the large shift in *V*_1/2_ caused by S182K. The slow time course of S182K currents resulted from specific increases in both the τ_slow_ and fraction of the slow component (Fig. 1H: τ_slow_ and reduction in *f*_fast_), but not the fast-kinetic component (Fig. 1H: τ_fast_). The specific effect on the slow component of the current kinetics hints that S182K may selectively stabilize the AO state. Overall, the mutational scan provided functional evidence that the CaM- and PIP_2_-interacting regions of the KCNQ1 S2-S3 linker, as seen in the cryo-EM structures, functionally impacts KCNQ1 activation.

### The S2-S3 linker interacts with CaM in KCNQ1 voltage-dependent gating

Although the S2-S3 linker residue S182 is located close to CaM in the CTD-bent structure (Fig. 2A), it is unclear whether this interface occurs under physiological conditions as the structure was solved without PIP_2_ (14, 15). Then, was the current effect induced by the mutation S182K (Fig. 1E-F) truly due to disruption of CaM-VSD interactions? We next examined whether these CaM-VSD interactions occurs during physiological gating, when PIP_2_ is available in the membrane. We undertook a second mutational scan of CaM residues close to the S2-S3 linker within the CTD-bent conformation (Fig. 2A-B). Each CaM mutant was co-expressed with WT human KCNQ1 channels and characterized in terms of *V*_1/2_. We found co-expression of CaM mutants at several positions (N98, Y100, N138, E140, E141) induced significant depolarizing shifts in the *V*_1/2_ (Fig. 2B, Supplementary Table 2). Similar to Q1-S182 mutations (Fig. 1E), mutations at CaM-Y100 demonstrated charge-dependent effects. CaM-Y100D induced a significant depolarizing shift in the *V*_1/2_, while CaM-Y100K yielded no change in the *V*_1/2_ (Fig. 2B-D). Additionally, CaM-Y100D altered current activation kinetics by increasing the fraction of the slow component current compared to WT (Fig. 2E, 2F: black to purple, reduction in *f*_fast_), but had no effects on τ_slow_ (Fig. 2F). This specific effect on the fraction of slow current component is similar to that seen with Q1-S182K mutation (Fig. 1H), and further suggests that CaM-Y100D may selectively stabilize the AO state. Altogether, the mutational scan revealed that CaM residues close to the S2-S3 linker within the CTD-bent conformation also affect KCNQ1 function.

**Figure 2.**
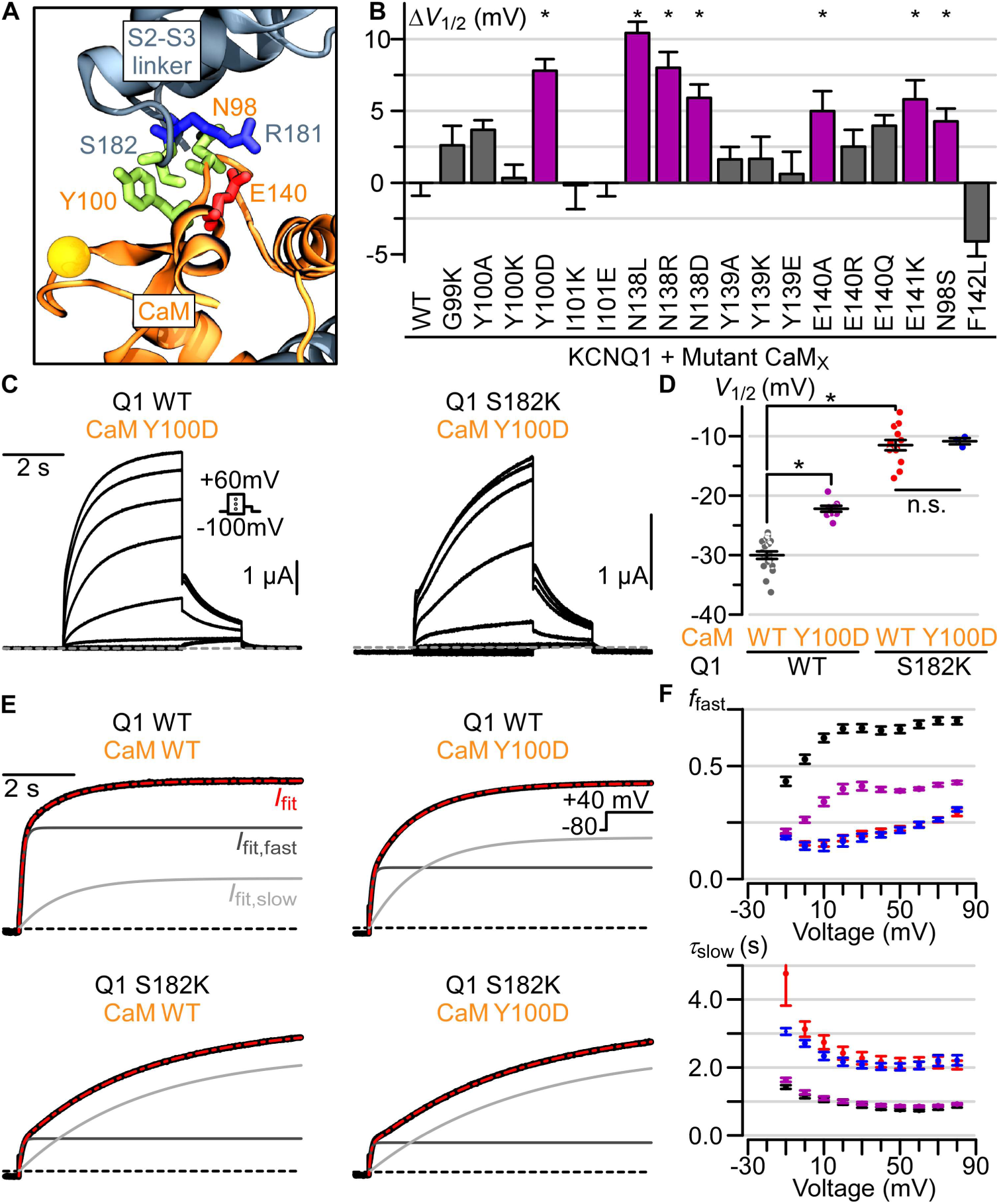
CaM interacts with the KCNQ1 S2-S3 linker during voltage-dependent gating. **(A)** Structural depiction of CaM residues predicted to interact with Q1-R181 and Q1-S182 in the CTD-bent structure. Residues depicted with sticks are colored according to residue type: positively (negatively) charged - blue (red), polar - green. Ca^2+^-ions are shown as yellow spheres. **(B)** Summary Δ*V*_1/2_ of human KCNQ1 co-expressed with mutant CaM. CaM-N98S and CaM-F142L are LQTS-associated CaM mutants. Purple bars and stars indicate significant Δ*V*_1/2_ from WT (Supplementary table 2). Error bars are SEM. **(C)** Exemplar currents from co-expression of CaM mutant Y100D with KCNQ1 WT (left) or KCNQ1 S182K (right) channels. Voltage protocol in inset. **(D)** Average *V*_1/2_ for KCNQ1 WT and S182K channels associated with either CaM WT or Y100D. Long bar: mean, short bars: SEM. Star: significantly different. P-values, calculated by t-test, are 2.18e-8 (Q1-WT/CaM-WT to Q1-WT/CaM-Y100D), 5.25e-17 (Q1-WT/CaM-WT to Q1-S182K/CaM-WT), and 0.73 (Q1-S182K/CaM-WT to Q1-S182K/CaM-Y100D). **(E)** Exemplar bi-exponential fits of activation kinetics for KCNQ1 WT and S182K channels associated with either CaM WT or Y100D. Black curve - ionic current; red curve - overall bi-exponential current fit (weighted sum of fast and slow components); light grey - slow component of the current fit; dark grey - fast component of the current fit; black dash line: 0 current. **(F)** Average population data for f_*fast*_ and τ_*slow*_ for KCNQ1 WT + CaM WT (black, *n* = 10), KCNQ1 WT + CaM Y100D (purple, *n* = 9), KCNQ1 S182K + CaM WT (red, *n* = 10), and KCNQ1 S182K + CaM Y100D (blue, *n* = 3). Error bars are SEM.

We next tested whether the S2-S3 linker and CaM interfaces interact during channel gating by combining KCNQ1 and CaM mutants. Because CaM-Y100 and Q1-S182 residues featured similar alteration to current phenotype, we co-expressed CaM-Y100D and Q1-S182K to test for interaction between the two residues. Although CaM-Y100D induced a depolarizing shift in the Q1-WT channel *V*_1/2_, CaM-Y100D failed to cause a depolarizing shift in the *V*_1/2_ of Q1-S182K channel (Fig. 2C-D). Likewise, while CaM-Y100D increased the slow component of the Q1-WT current (a reduction in *f*_fast_), CaM-Y100D did not induce any further enhancement in the slow component of Q1-S182K channels (Fig. 2E-F). These findings indicate that the single mutation Q1-S182K confers the full functional impact of the double mutations Q1-S182K and CaM-Y100D. Given that Q1-S182 and CaM-Y100 are in proximity within the CTD-bent structure, the non-additive shifts of mutant *V*_1/2_ are consistent with a functional interaction between Q1-S182 and CaM-Y100 during KCNQ1 activation. By contrast, CaM-Y100D induced a significant depolarizing shift in the Q1-R181E channel *V*_1/2_ (Supplementary Fig. 1A), indicating that Q1-R181 and CaM-Y100 may affect channel gating by different mechanisms. Taken together, these data suggest a specific Q1-S182 and CaM-Y100 interaction during KCNQ1 gating, providing evidence that the mutation effects at Q1-S182 and CaM-Y100 are due to disrupted CaM-VSD interactions. This finding also shows that CaM-VSD interactions observed in the CTD-bent conformation indeed occur during the physiological KCNQ1 activation process despite the presence of PIP_2_ in the membrane.

To further investigate the interactions at the CaM-VSD interface, we performed molecular dynamics (MD) simulations of KCNQ1/CaM movements at physiological temperature. Statistics calculations from the multiple frames in the simulations allowed modeling of dynamic interactions among residues that are not revealed by the structure itself. We analyzed MD simulations of the human and *Xenopus* KCNQ1/CaM cryo-EM structures (14, 15) in the CTD-bent conformation. All MD simulations ran for aggregated time of at least 1 µs (Supplementary Fig. 2A-B). Potential interactions between CaM and the S2-S3 linker were examined by computing the minimum distance between non-hydrogen atoms of pairwise CaM-VSD residues in each simulation frame. The MD simulations of both the human and Xenopus KCNQ1/CaM structures revealed interaction distances consistent with salt-bridge/hydrogen bond interactions (∼3.5 Å) between the pairs Q1-S182/CaM-Y100 and Q1-S182/CaM-N138 (Supplementary Fig. 3). We also found close interaction distances (∼5 Å) between Q1-S182 and CaM-N98 (Supplementary Fig. 3). Our results therefore showed that the CaM-VSD interactions seen in the CTD-bent structure remain intact after relaxation by MD simulations. Notably, these key residues at the CaM-VSD interface found in the simulations are consistent with the mutagenesis scan (Figs. 1-2). We also identified a salt-bridge between residues Q1-R181 and CaM-E140 in the simulations (Supplementary Fig. 3). However, co-expression of CaM-E140 mutants with the WT KCNQ1 channel showed only modest effects (Fig. 2B), rendering functional detection and validation of Q1-R181/CaM-E140 interaction difficult. Despite this, the MD simulations generally agreed with the mutagenesis results and provided further support that the CaM-VSD interactions seen in the CTD-bent conformation are stable. Taken together, the combined functional and simulation results suggest CaM interacts with the KCNQ1 S2-S3 linker during gating, even under conditions when PIP_2_ is available.

The results so far provide a preliminary hypothesis to contextualize the two KCNQ1/CaM cryo-EM structural states within the voltage-dependent gating process. We found that disruption of the CaM-VSD interface by Q1-S182K and CaM-Y100D enhanced the slow component of current activation but had no effect on the fast kinetic component (Fig. 1G-H, Fig. 2E), suggesting that the AO state is promoted by the loss of CaM-VSD interactions. As a major feature of the CTD-straight conformation is the lack of CaM-VSD interactions, along with the fact that the CTD-straight structure features the VSD at the activated state and the pore dilated (14, 15, 21), our results provide experimental evidence to support the idea that the CTD-straight structure corresponds to the functional AO state. On the other hand, our finding that CaM-VSD interactions occur during activation suggests that the CTD-bent structure also corresponds to a functional state in the gating pathway, although the structure was determined in the absence of PIP_2_ (14). How can the CTD-bent and the CTD-straight conformations both occur during gating, when they feature seemingly mutually exclusive CaM-VSD and PIP_2_-VSD interactions? A likely explanation is that CaM binding and PIP_2_ binding to the S2-S3 linker are state-dependent, such that KCNQ1 transitions between the two conformations during activation. In this hypothesis, the exchange of S2-S3 linker binding partner (CaM or PIP_2_) during activation may facilitate transitions between the gating states associated with the two structures.

### The CTD-straight conformation features properties specific for the AO state

To further understand the role of the CaM-VSD interactions for transitioning between the CTD-bent and CTD-straight conformations, and also to further define the gating states associated with these conformations, we turned to MD simulations and network analysis. In KCNQ1 gating, the movements of the VSD S4 gating charges are sensed by nearby interacting residues and transmitted to the pore domain via a chain of residue interactions. This chain of interactions effectively constitutes VSD-pore allosteric, or coupling, pathways for the channel to open. The CaM-VSD interactions are present in the CTD-bent structure but absent in the CTD-straight structure (14, 15) (Figs. 1A-B). To investigate how this change in the CaM-VSD interactions affects VSD-pore allostery, we employed MD simulations to track all residue interactions within both structures, followed by network analysis to identify VSD-pore allosteric pathways in the two conformations (Fig. 3A). Consequently, this enabled us to contrast the specifics of how the CaM-VSD interactions influence KCNQ1 VSD-pore allostery.

**Figure 3.**
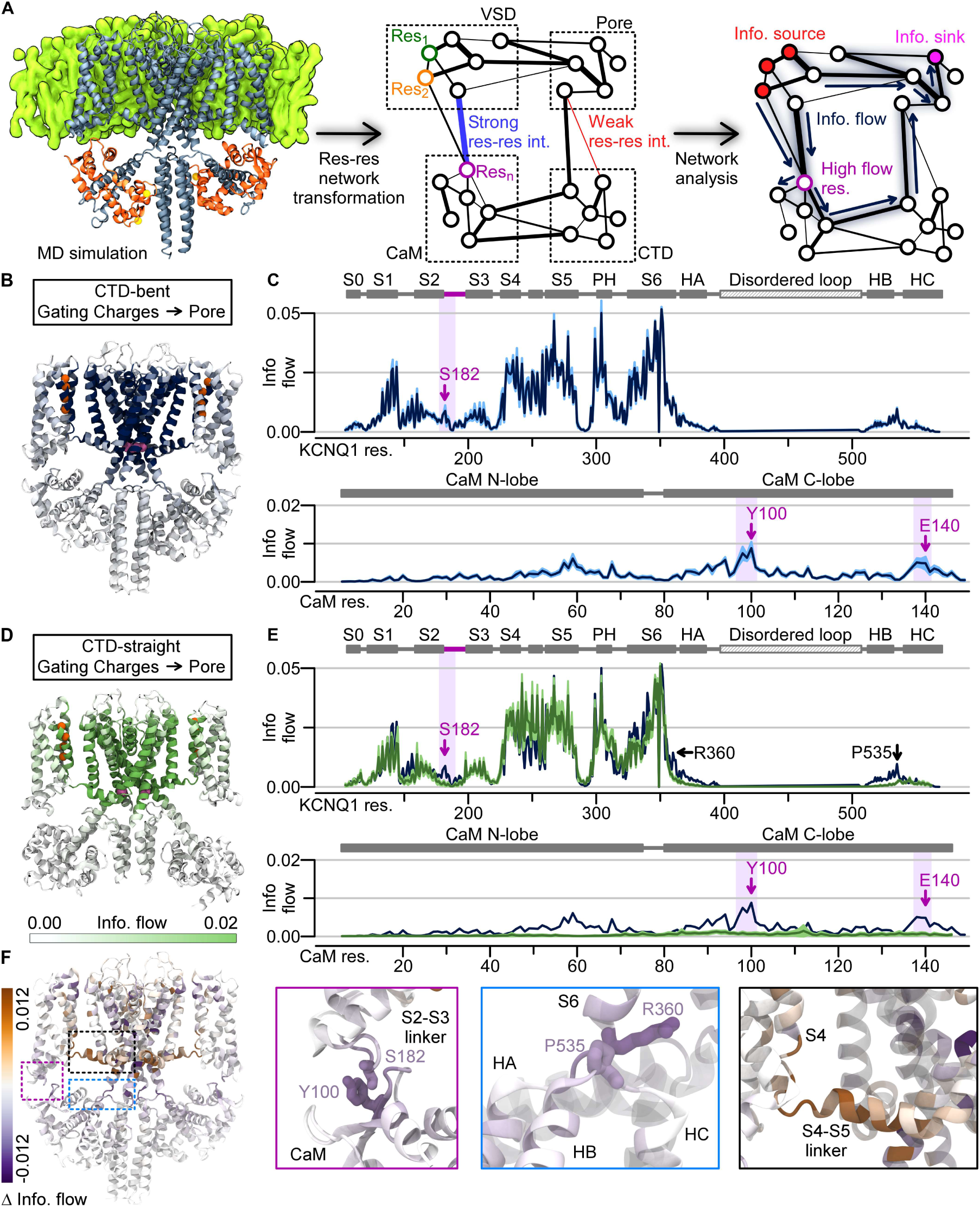
Network analysis of molecular dynamics simulations demonstrates that the CaM-VSD interactions modulate KCNQ1 VSD-pore allosteric pathways. **(A)** Flowchart of MD simulation analysis pipeline. The KCNQ1/CaM complex cryo-EM structure was inserted into a membrane and simulated with MD in explicit solvent. The protein was transformed into a network; each node is a residue while each edge weight corresponds to interaction strength between residues. Network analysis was applied to track information flow from defined source residues (nodes) to sink residues. High-information-flow nodes carry key allosteric pathways from the source to sink residues. **(B)** CTD-bent information flow (VSD gating charges to the pore) through each residue projected with color intensity onto the KCNQ1/CaM CTD-bent structure. Dark color - high current flow (capped at 0.02). Source residues (gating charges) - orange spheres, sink residues (pore) - magenta spheres. Two full subunits out of four are shown for clarity; the VSDs, helices A/B and CaMs in the front and back subunits are omitted. **(C)** Information flow profiles of human KCNQ1 (dark blue) and CaM (black) along residue sequence. Light purple shadings: rough position of the S2-S3 linker/CaM interface. Purple arrows: peaks in the S2-S3 linker/CaM interface. Protein-specific domains and helices are labelled above the plots: PH = pore helix; HA/B/C = helix A/B/C. **(D)** Structural depiction of the information flow in the CTD-straight structure, similar to panel B. **(E)** Information flow profile of CTD-straight conformation along the KCNQ1 sequence (green). Dark blue: same as in panel **(C)**. Green shading: the CTD hub. **(F)** Left: Δinformation flow projected on the structure (CTD-straight versus CTD-bent information flow). Right: detailed views of the main interfaces exhibiting large increase (brown) or decrease (purple) in information flow.

Network analysis is an approach which previously has shown effective in protein allostery identification (37–41). First, the MD trajectories of the KCNQ1/CaM complex were converted into a residue interaction network representation (Fig. 3A). In this network, each node corresponded to an individual amino acid residue within the KCNQ1/CaM complex. The weights on the connections (edges) between nodes capture interaction strength between the residues, which were defined by spatial proximity and correlated residue movements within the MD trajectories. The final network thus encodes all residues (nodes) and interactions (edges) within the KCNQ1/CaM complex. Next, we extracted allosteric pathways between the KCNQ1 VSD and the pore by measuring how information flows through the network. We defined the VSD gating charges as the “source”, from which information originates, and tracked how information flows to arrive at the “sink”, the pore-gate residue S349 (Fig. 3A, 3B: orange and magenta spheres, respectively). The underlying idea is that a perturbation of residue interactions, such as those induced by the movements of S4 gating charges (the source), spreads to other residues (nodes) via diffusion along the KCNQ1/CaM complex (network) before they arrive at the pore (the sink). We employ the method most commonly referred to as “current flow betweenness” (38, 42, 43) to account for all pathways from the source to the sink. Here, we refer to this method as “information flow analysis” to avoid confusion with ion channel terminologies. Critically, the ability of one residue to transmit information to another depends on the edge weight, which encodes the strength of their interaction. The flow across an edge in opposing directions are cancelled out, leaving the net positive information flow. Thus, key residues and pathways that participate in productive information transmission from the gating charges to the pore gate are systematically highlighted by strong information flow. This information flow analysis is carried by the dynamic interactions among all residues such that it is not readily shown by the structural data that only capture one or a few frames of the dynamic motion of the protein. However, we emphasize that this analysis, based on structural data, identifies allosteric pathways specific to the simulated channel state.

We applied network analysis on MD simulations of the human KCNQ1/CaM CTD-bent and CTD-straight structures (Fig. 3B-E, Supplementary Fig. 2C). To visualize the predicted spatial pattern of major VSD-pore allosteric pathways within each state, we projected the calculated information flow at each residue onto the respective structures (Fig. 3B,3D), such that darker color signifies higher flow. Figure 3C and 3E display the information flow along the KCNQ1 and CaM residue sequences. As expected, the highest information flow residues in both conformations were observed at the S4-S5 linker/S6 and the S4/S5 interfaces within the transmembrane domains. These two interfaces are well-studied in domain-swapped K_V_ channels as critical for VSD-pore coupling (40, 44, 45), including KCNQ1 (18). The network analysis thus recapitulated the importance of these transmembrane motifs in KCNQ1 VSD-pore coupling. We also performed the same analysis on simulations of the *Xenopus* KCNQ1/CaM CTD-bent structure. The results were close to identical to the analysis of the human KCNQ1/CaM complex (Supplementary Fig. 4A-D), reinforcing the findings from the human KCNQ1/CaM simulations.

To examine how the switch from the S2-S3 linker binding CaM to binding PIP_2_ influences VSD-pore allosteric pathways, we calculated the difference between the CTD-bent and CTD-straight information profiles. The resulting change in information flow (Δinformation flow) is projected onto the cryo-EM structure as shown in Figure 3F, with brown and purple indicating enhanced and reduced flow strength in the CTD-straight over the CTD-bent conformation. This revealed flow strength differences at distinct locations due to the transition from the CTD-bent to the CTD-straight conformation.

First, the CTD-straight conformation exhibited a clear reduction in information flow at the CaM/VSD interface compared to the CTD-bent conformation (Fig. 3F, purple box). The CTD-bent flow profile featured clear peaks at the S2-S3 linker and CaM (Fig. 3B-C, purple shading), while the CTD-straight flow profile lacked the same peaks (Fig. 3D-E, Supplementary Fig. 4E). This loss of the CaM/VSD peaks is associated with the loss of CaM-VSD interactions, consistent with the notion that the S2-S3 linker/CaM peaks in the CTD-bent flow profile are attributed to CaM-VSD interactions. Notably, the residues corresponding to the S2-S3 linker/CaM flow peaks were Q1-S182, Q1-R181, CaM-N98, CaM-Y100, and CaM-E140 (Fig. 3C), consistent with our mutational scans (Figs. 1-2). The S2-S3 linker peaks are weaker compared to the strongest peaks at the classic transmembrane interfaces, suggesting that the coupling pathways through the CaM-VSD interface are less critical than those via the transmembrane interfaces. Nevertheless, these results show that the CaM-VSD interactions play a role, albeit small, in the VSD-pore allostery pathways in the CTD-bent conformation. This pathway is lost upon the transition to the CTD-straight conformation.

Beyond the CaM/VSD interface, the CTD also exhibited reduced flow strengths from the CTD-bent conformation to the CTD-straight conformation. The CTD region with the most reduced flow strengths involved the S6-HA and HB-HC linkers (Figure 3F, blue box). The concurrent flow reductions at the CaM/VSD interface and the CTD hint that these two regions might be linked allosterically and work together to affect KCNQ1 pore opening. This phenomenon will be further explored later in this study.

In contrast to the reduced flow strengths at the CaM/VSD interface, the CTD-straight conformation interestingly featured enhanced information flow in C-terminus of S4 and N-terminus of the S4-S5 linker (Figure 3F, black box). This result correlates with prior functional and simulation studies to further supports the suggestion that the CTD-straight structure corresponds to the AO state. We previously found that KCNQ1 VSD activation triggers a two-stage VSD-pore coupling mechanism, such that two sets of VSD-pore interactions induce channel opening with the VSD movements to the intermediate state and the activated state (18). Particularly, the VSD-pore coupling interactions specific for the AO state involve the C-terminus of S4, the N-terminus of the S4-S5 linker, and part of S5 and S6 (18). These critical coupling motifs for the AO state are closely correlated with the locations featuring enhanced information flow in the CTD-straight conformation (Fig. 3F, black box). The CTD-straight conformation thus highlights the structural basis of the VSD-pore coupling of the AO state, providing another line of evidence that the CTD-straight conformation corresponds to the functional AO state. Importantly, the flow strength enhancement at these AO coupling motifs is accompanied by the loss of CaM-VSD interface, suggesting that the loss of CaM-VSD interactions reinforces AO-state coupling interactions.

Taken together, the comparative network analysis provided two insights on the role of CaM-VSD interactions in KCNQ1 VSD-pore coupling. First, the presence of CaM-VSD interface in the CTD-bent state provides a minor VSD-pore coupling pathway (Fig. 3B-E). Second, the loss of CaM-VSD interactions in the CTD-straight state enhances AO-state specific coupling interactions in the channel transmembrane domains (Fig. 3F). This result suggests that the CaM-VSD interactions play a role in reinforcing key AO state coupling interactions during the transition from the CTD-bent to CTD-straight conformation, thereby facilitating channel transition into the AO state.

### CaM-VSD interactions control the transition to the AO state

So far, our mutational scan data suggested that KCNQ1 may transition between the CTD-bent and CTD-straight conformations during activation. Furthermore, our MD network analysis predicted that the switch from the S2-S3 linker binding CaM to PIP_2_ may promote channel transition into the AO state. To test this hypothesis, we probed how disrupting CaM-VSD and PIP_2_-VSD interactions affect distinct aspects of KCNQ1 gating mechanism using voltage-clamp fluorometry (VCF). The KCNQ1 VSDs are fluorescently labelled in VCF to enable simultaneous monitoring of VSD movements and pore opening during activation. We specifically applied VCF on the Q1-R181E and Q1-S182K mutants, which disrupt the S2-S3 linker interaction with PIP_2_ and CaM, respectively.

Figure 4A illustrates VCF recordings from KCNQ1 pseudoWT (psWT) channels, which carries three mutations (C214A/G219C/C331A, hereby denoted Q1*) to enable fluorescent labeling specifically to G219C. This Q1* psWT channel construct has been extensively utilized and well-validated in prior studies across many laboratories as the WT control for KCNQ1 VCF studies (8, 17–19, 36). The Q1*-psWT channels featured robust changes in fluorescence signal upon membrane depolarization corresponding to VSD activation, which tracks closely with ionic currents correlating with pore opening (Fig. 4A, blue and black exemplars). Next, we applied VCF to the Q1*-R181E mutant to assay how disrupted PIP_2_ binding to the S2-S3 linker affects VSD activation. We found a significant depolarizing shift in the voltage dependence of VSD activation of Q1*-R181E (F-V, Supplementary Fig. 1B-C, F_1_ curve). This result draws a connection between PIP_2_ binding to the S2-S3 linker and VSD activation movement. If PIP_2_ binding is state-dependent for when the VSD is activated, as suggested by the CTD-straight structure (15), then PIP_2_ binding also facilitates VSD activation. The disruption of PIP_2_-VSD interactions by the Q1-R181E mutation can thus induce depolarizing shifts in VSD activation. This state-dependent PIP_2_-VSD interaction is consistent with the idea that PIP_2_ competes with CaM at the S2-S3 linker upon VSD movement into the activated state during gating, leading to a state-dependent loss of CaM-VSD interactions.

**Figure 4.**
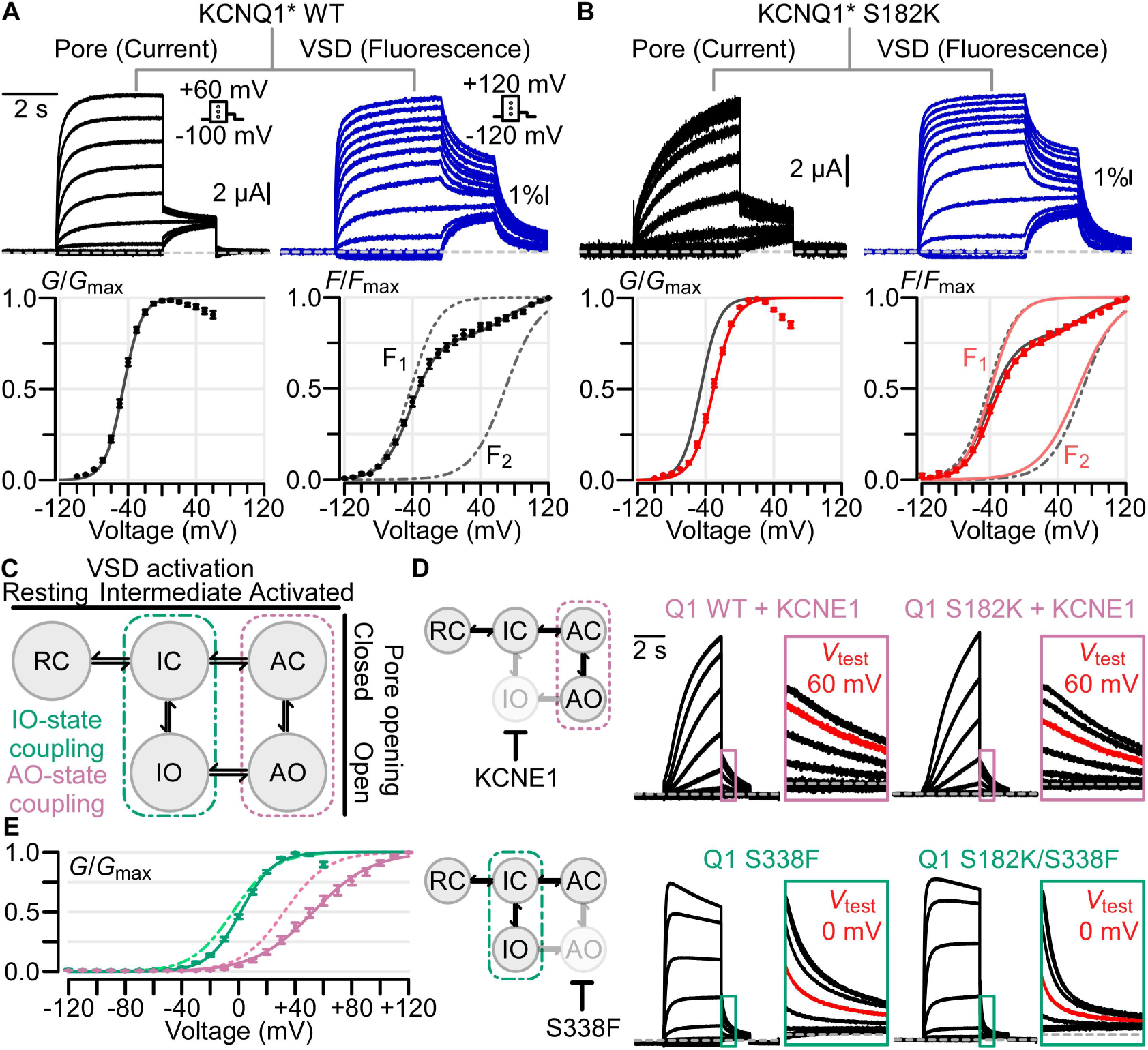
CaM-VSD interactions control the transition to the AO state. **A)** Voltage-clamp fluorometry (VCF) experiments using KCNQ1* psWT. Left and right columns: exemplar ionic current/VSD fluorescence signal measurements (Voltage protocols in inset) and average G-V/F-V relationships, respectively. The F-V relationship (solid black curve) is fitted with a double Boltzmann equation (F_1_ and F_2_: dotted black curves), see Materials and Methods. **(B)** Same as panel A, but for KCNQ1* S182K mutant channel. In G-V and F-V plots: black - psWT; red - S182K. **(C)** KCNQ1 gating model with two open states, IO and AO. The KCNQ1 VSD transitions into the intermediate and activated conformations engage distinct sets of VSD-pore coupling interactions (green and violet boxes) to trigger pore opening. **(D)** Left: cartoon schematic illustrating the effect of KCNE1 co-expression (top) and Q1-S338F mutation (bottom) to ablate the IO and AO states respectively to leave the KCNQ1 to conduct only at one open state. Right: current exemplars for KCNQ1 WT and mutant channels designed to probe whether Q1-S182K mutation specifically affects IO or AO state. Each box shows enlarged tail currents, with the red trace indicating the tail current associated with the labeled test voltage, which shows the shift of voltage dependence when comparing the WT and mutant. Voltage protocol same as in Panel A, except for exemplar currents with KCNE1 co-expression test pulses range from −120 mV to +100 mV. **(E)** Average G-V curve and Boltzmann fits for Q1-WT+KCNE1 (dotted pink, n = 5), Q1-S182K+KCNE1 (solid pink, n = 7), Q1-S338F (dotted green, n = 6), Q1-S182K/S338F (solid green, n = 11).

We next studied the Q1*-S182K mutant to measure how disruption of CaM-VSD interactions affects VSD activation. VCF recordings of the Q1*-S182K channels showed a depolarizing shift in G-V and slowdown in current activation kinetics (Fig. 4B). However, little change in the voltage dependence or kinetics of VSD activation was observed by the mutation (Fig. 4B). This finding indicates that the disruption of CaM-VSD interactions did not directly modulate VSD activation. Therefore, the mutation changed pore openings by altering either the VSD-pore coupling or the pore directly. Since the mutation is in the S2-S3 linker, spatially far from the pore domain, it is likely that the mutation altered VSD-pore coupling. This result is consistent with the suggestion that the disruption of CaM-VSD interactions facilitates channel transition to the AO state by enhancing VSD-pore coupling pathways specifically important for the AO state (18) (Fig. 3F).

We next asked if the role of CaM-VSD interactions is truly specific to modulating transition into the AO state. To answer this question, we studied how CaM-VSD interactions affect the IO versus the AO state. We disrupted the CaM-VSD interactions with Q1-S182K mutation in modified KCNQ1 channels that selectively open only to the AO or IO state. Our previous work showed that co-expression of the auxiliary subunit KCNE1 with KCNQ1 suppresses the IO state (8, 18, 21), forcing KCNQ1 to conduct only in the AO state (Fig. 4D, top). Conversely, the KCNQ1 mutation S338F selectively abolishes the AO state by disabling the VSD-pore coupling when the VSD is at the activated state, thus leaving the channel to only conduct at the IO state (Fig. 4D, bottom) (18, 23, 24). We first tested how disrupting CaM-VSD interactions affect the AO state by KCNE1 co-expression with Q1-S182K channels. The G-V relationships (Fig. 4D-E) showed that Q1-S182K+KCNE1 maintained a 20-mV right shift in *V*_1/2_ compared to Q1-WT+KCNE1, consistent with the shift observed between Q1-S182K and -WT channels without KCNE1 (Fig. 1E). This data provides another line of evidence that the CaM/VSD interactions modulate channel transition into the AO state. We then probed whether CaM-VSD interactions modulate the IO state with the Q1-S182K/S338F double mutant. Distinct from the KCNE1 co-expression experiments, the Q1-S182K mutation failed to induce a depolarizing shift in the *V*_1/2_ compared to Q1-S338F (Fig. 4D-E). Introduction of the Q1-S182K mutation also failed to cause a slowdown of Q1-S338F current activation kinetics. These results indicate that CaM-VSD interactions exert no modulation on the IO state. This lack of IO state modulation is also consistent with the lack of S182K effect on the KCNQ1 fast-kinetic current component (Fig. 1H), which approximates the IO state. Together, these results show that the CaM-VSD interactions exert a selective modulation of the KCNQ1 AO state. Along with the finding of state-dependent PIP_2_-VSD interaction, all these results suggest that the loss of CaM-VSD interactions during KCNQ1 gating, as driven by PIP_2_ competition, selectively controls channel transition into the AO state.

### A hub in the KCNQ1 C-terminal domain cooperates with the CaM-VSD interactions to control the transition to the AO state

Does the CaM-VSD interactions work alone or with other structural motifs to control KCNQ1 transition into the AO state? Our MD network analysis identified a reduction in information flow strength in the CTD that accompanied the loss of CaM-VSD interactions (Fig. 3F), hinting the CTD might be allosterically downstream of CaM-VSD interactions. Moreover, the cryo-EM structures showed that the loss of CaM-VSD interactions was accompanied by major conformational rearrangements in the CTD (Fig. 1A-B). Specifically, the “RQKH” motif within the KCNQ1 S6-HA linker was shown to undergo a loop-to-helix transformation from the CTD-bent to the CTD-straight conformation (15). These data together hint that the CaM-VSD interactions may work together with the CTD to affect channel transition into the AO state. We thus further pursued this hypothesis and turned back to MD network analysis to systematically identify key residues within the CTD which work together with the CaM-VSD interface.

To identify CTD structural motifs which might be allosterically downstream of the CaM-VSD interaction, we performed an alternate network analysis on the simulations of the CTD-bent conformation. The information flow in the CTD-bent conformation was computed using all residues within the CaM C-lobe, as opposed to the S4 gating charges, as source residues (Fig. 5A). As described previously, the information profile calculated with the S4 gating charges as the source corresponds to VSD-pore allosteric pathways in the CTD-bent structure. By contrast, the profile calculated with the CaM C-lobe as the source revealed how perturbation in CaM is propagated to the pore domain residue Q1-S349. Thus, residues which are specifically sensitive to changes in CaM, such as the loss of CaM-VSD interactions, should feature higher information flow when CaM is defined as the source versus the gating charges. The Δinformation flow between the CaM C-lobe and gating charge information profiles is projected onto the CTD-bent structure in Figure 5B. When using CaM C-lobe as the source, there was a general reduction in flow strengths through the transmembrane domains but a general enhancement of flow strengths through the CTD (Fig. 5A-B, Supplementary Fig. 5A). This indicates that perturbation in CaM is preferentially propagated to the pore through the CTD, consistent with the idea that loss of CaM-VSD interactions might work with the CTD to affect the pore. The CTD residues featuring the greatest flow enhancement included Q1-R360 (S6-HA linker) and Q1-P535 (HB-HC linker) (Supplementary Fig. 5A; green shading). Similar results were obtained with the same analysis on the Xenopus KCNQ1/CaM complex (Supplementary Fig. 5B-C), further validating the conserved importance of Q1-R360 and Q1-P535. Moreover, in our previous Δinformation flow comparison between the CTD-bent to CTD-straight conformation, these same two CTD residues (Q1-R360, Q1-P535) exhibited simultaneous reduction in information flow with the CaM/VSD interface (Fig. 3F, black box). Taken together, these results suggest that residues Q1-R360 and Q1-P535 are allosterically coupled to the CaM-VSD interactions. Consequently, the network analysis predicted that these CTD residues are important in conjunction with the CaM-VSD interactions to affect channel transition into the AO state.

**Figure 5.**
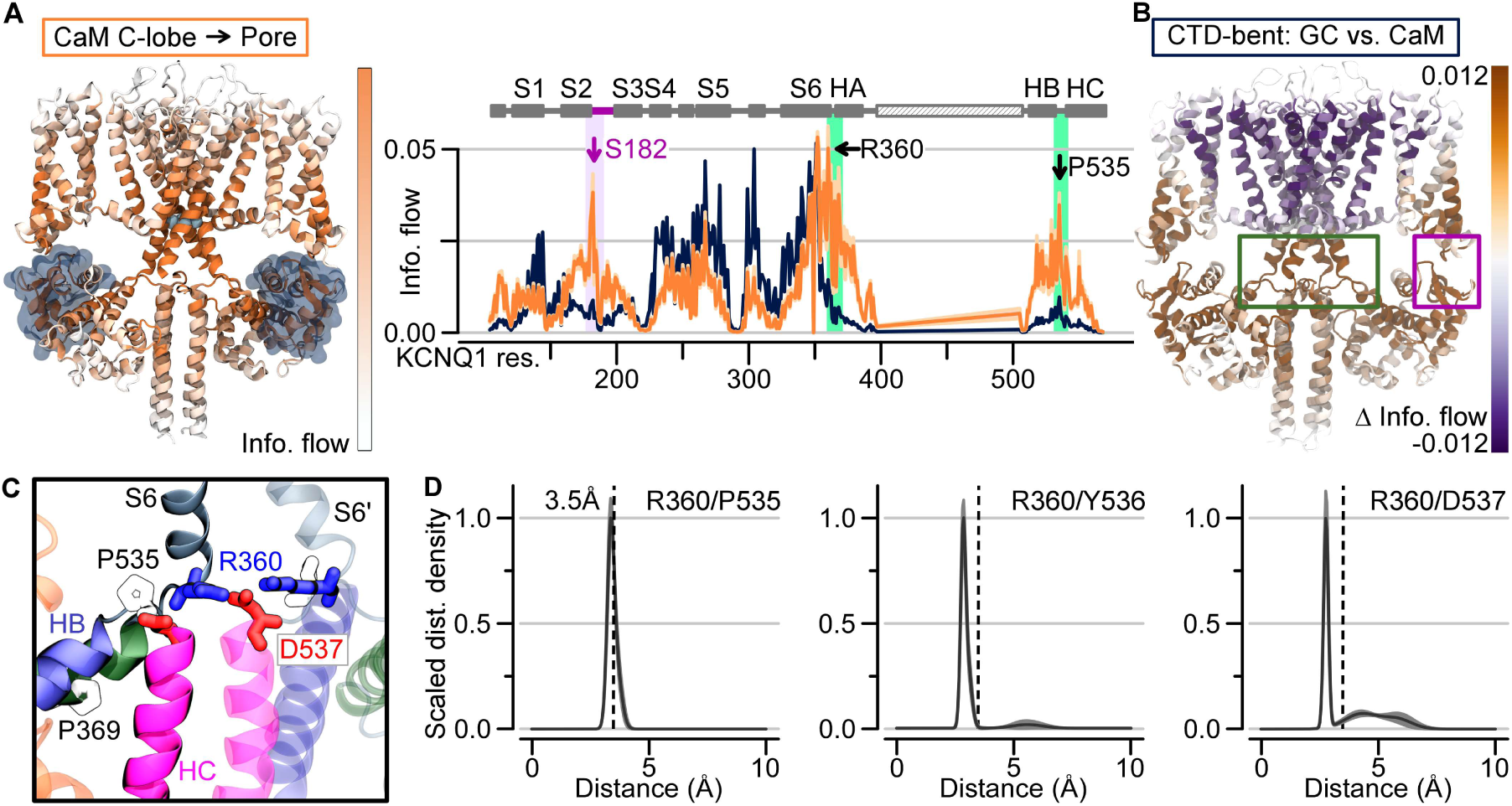
A hub in the KCNQ1 C-terminal domain cooperates with the CaM-VSD interactions to control the transition to the AO state. **(A)** Left: Information flow strengths projection onto the CTD-bent structure, capped at 0.02. For clarity, VSDs, helices A/B and CaMs in the front and back subunits are omitted. Right: Orange: Information flow profile along KCNQ1 sequence (CaM C-lobe used as the source) in the CTD-bent structure; Dark blue: Information flow profile with gating charges as source. Purple shading: information flow peak in the S2-S3 linker/CaM interface. Green shading: peak between the helix B/C and helix A/S6 loops. **(B)** Δinformation flow projected on the CTD-bent structure (CaM C-lobe versus the gating charges as the source). Dark violet residues: decreased information flow as a result of using CaM C-lobe as source; brown: increased information flow. **(C)** Structural depiction of residues in the CTD hub. Green ribbons - HA; green ribbons - HB; pink ribbons - HC; grey ribbons - S6. S6’ is from a neighboring subunit. Residues depicted with sticks are colored according to residue type: positively (negatively) charged - blue (red) and hydrophobic - white. **(D)** Scaled distance density of interacting residues in the CTD hub.

Does the spatial arrangement of Q1-R360 (S6-HA linker) and Q1-P535 (HC-HC linker) support the hypothesis that these CTD residues are allosterically downstream of the CaM-VSD interactions? Although these residues are far apart in terms of primary sequence, we found a cluster of residues, including Q1-R360 and Q1-P535, which formed a structural motif that pinches the S6-HA and HB-HC linkers together in the CTD-bent simulations (Fig. 5C). These interactions included a close interaction between Q1-P535 and Q1-R360, a salt-bridge between Q1-D537 and Q1-R360, as well as hydrogen bonding between Q1-Y536 and Q1-R360 (Fig. 5D). Notably, these residues and interactions are conserved between the human and *Xenopus* KCNQ1 channels (Supplementary Fig. 5D,E). Interestingly, Q1-R360 interacts with Q1-Y536 and Q1-D537 of the neighboring subunit (i.e. inter-subunit) in the CTD-bent structure, but these interactions switch to intra-subunit in the CTD-straight structure. The residue Q1-R360 further corresponds to the arginine in the “RQKH” motif that undergoes a loop-to-helix transition between the CTD-bent and CTD-straight structures (15). Overall, these findings are consistent with the idea that this cluster of residues may form a critical hub (“CTD hub”) that facilitates CTD conformational rearrangement. Our analysis further predicted that this CTD hub allosterically works together with the CaM-VSD interface to control the channel transition into the AO state.

### Functional validation that the CTD hub cooperates with CaM-VSD interactions in KCNQ1 gating

To test the prediction that the CTD hub is allosterically downstream of the CaM-VSD interactions to modulate KCNQ1 transition into the AO state, we performed functional electrophysiology experiments. Specifically, we examined the two key CTD residues identified by network analysis, Q1-R360 and Q1-P535. We additionally tested another HA residue, Q1-P369, which showed a slightly increased peak of information flow when CaM C-lobe was used as the source in the CTD-bent conformation (Fig. 5A). Notably, P369 is well conserved in the KCNQ family and mutation at the analogous position in KCNQ4 (P369R) is linked to intellectual disability (46). If the loss of CaM-VSD interactions and the CTD hub are allosterically coupled to control channel transition into the AO state, then double mutations at the S2-S3 linker and the CTD should feature non-additive changes to shift in *V*_1/2_, and likely non-additive effects on current kinetics. We thus characterized S2-S3 linker S182K and CTD double mutants to test this hypothesis (Fig. 6A).

**Figure 6.**
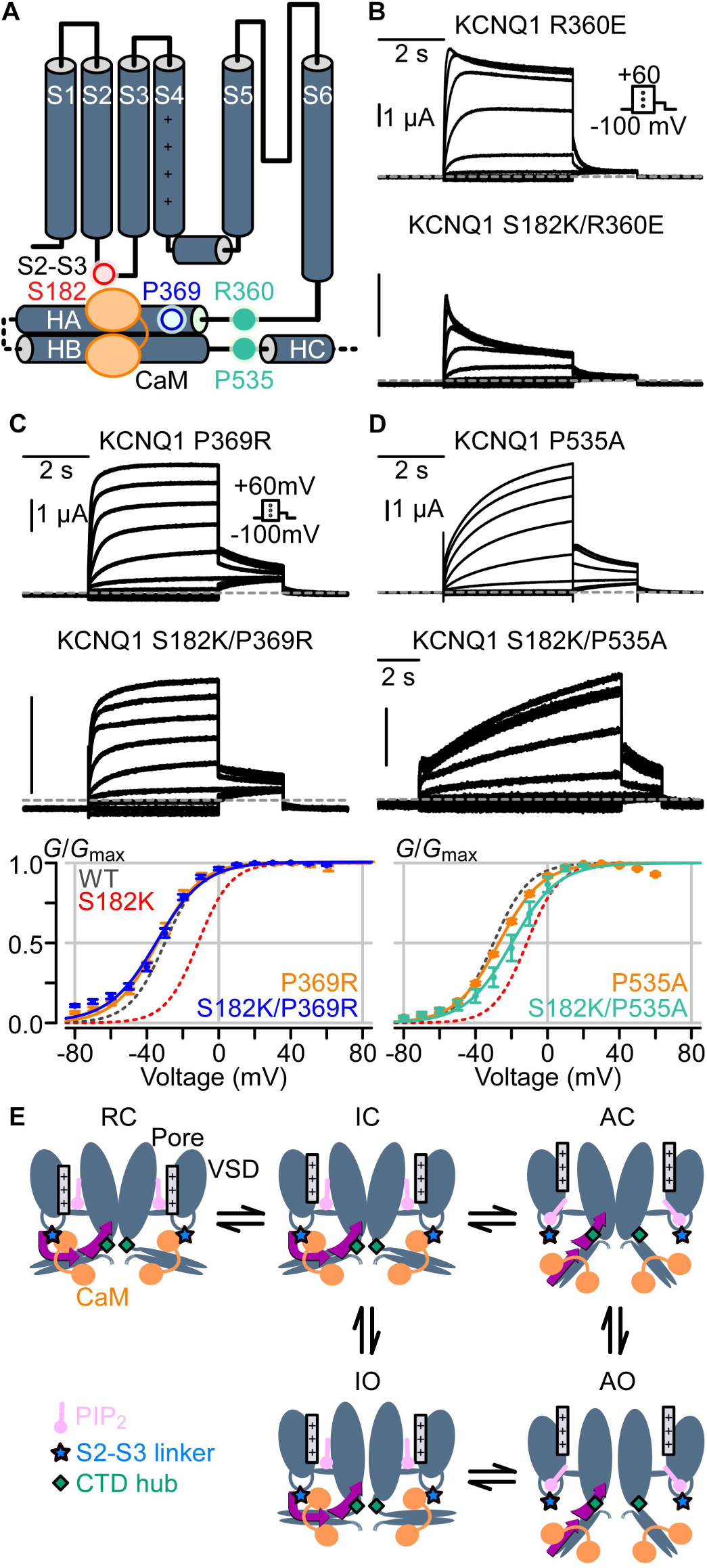
Functional validation that the CTD cooperates with CaM-VSD interactions in KCNQ1 gating. **(A)** Cartoon depiction of KCNQ1 channel with key CTD residues (blue and green circles) predicted to be allosterically downstream of the CaM-VSD interaction (red circle). **(B)** Current exemplars for mutants involving S2-S3 linker (S182K) and the S6-HA loop (R360E). Voltage protocol in inset. (C-D) Top and middle: current exemplars for mutants involving the S2-S3 linker (S182K), the proximal helix A (P369R), and the HB-HC loop (P535A). Bottom: average G-V relationships calculated for the respective mutants. Dashed lines are G-V fits for WT (grey) and KCNQ1-S182K (red). N = 7 (KCNQ1-P369R), 5 (KCNQ1-S182K/P369R), 8 (KCNQ1-P535A), 3 (KCNQ1-S182K/P535A). **(E)** KCNQ1 voltage-dependent gating model integrating CaM and PIP_2_. Color designations: KCNQ1 channel (dark gray), KCNQ1 S2-S3 linker (light blue star), KCNQ1 CTD hub (green diamond), CaM (orange), PIP_2_ (violet). Helix C and D are not drawn for clarity. PIP_2_ interaction with the KCNQ1 S2-S3 linker is dependent on the VSD state. PIP_2_ does not interact with the S2-S3 linker when the VSD is in the resting **(R)** or intermediate **(I)** states, thus CaM favorably interacts with the S2-S3 linker. Upon VSD transition into the activated **(A)** state, PIP_2_ favors interaction with the S2-S3 linker and competes with CaM. This competition disengages CaM from the S2-S3 linker and allows the CTD to undergo large conformational rearrangement pivoted at the CTD hub (green diamond) to induce the AO state.

We first probed the CTD S6-HA linker single mutation Q1-R360E, which yielded robust ionic currents with slight inactivation (Fig. 6B), consistent with prior studies (36). In contrast to the slowly activating Q1-S182K single mutant current, the double mutant Q1-S182K/R360E exhibited a rapid inactivation current phenotype (Fig. 6B). We did not calculate the *V*_1/2_ of Q1-S182K/R360E double mutant due to the strong current inactivation. Nevertheless, the dramatic switch in current activation kinetics from the single to double mutant channel is consistent with allosteric interaction between S182 and R360. Next, we found that the Helix A mutant P369R and the HB-HC linker mutant Q1-P535A both featured hyperpolarized *V*_1/2_ compared to the WT channels (Fig. 6C-D). Q1-P369R featured a rapid current activation kinetics (Fig. 6C), while Q1-P535A exhibited a markedly slowed activation kinetics (Fig. 6D). Interestingly, both double mutants Q1-S182K/P369R and Q1-S182K/P535A featured hyperpolarized *V*_1/2_ compared to Q1-S182K (Fig. 6C-D), reversing the *V*_1/2_ shift induced by the Q1-S182K mutation and demonstrating non-linear shifts in *V*_1/2_. Moreover, the double mutant Q1-S182K/P369 featured a rapidly activating current kinetics, completely reversing the slowly activating phenotype of the Q1-S182K single mutant. Taken together, we found residues in the KCNQ1 CTD (P369, P535) that interfered with ability of the CaM-VSD interface to affect transition into the AO state. These results provide experimental evidence that the loss of CaM-VSD interactions works in concert with the CTD hub to control channel transition into the AO state. As mentioned, these residues are dispersed in terms of primary sequence (S6-HA linker, helix A, and HB-HC linker). This indicates that a switch in CaM-VSD interactions extensively employs the tertiary structure of the CTD to control channel transition into the AO state.

## Discussion

With increasing number of resolved atomic structures of full-length ion channels, correlating these structures to their gating states is becoming a wide-spread question. In the KCNQ1/CaM structure determined without PIP_2_, which we called the CTD-bent conformation, the VSD is activated and the pore is closed (14, 15). The S6-HA linker bends to bring CaM near the VSD and enable CaM C-lobe interaction with the S2-S3 linker (14, 15, 21) (Fig. 1A). The KCNQ1/CaM structure solved with PIP_2_, which we termed CTD-straight conformation, features activated VSD and dilated pore (15, 21). Strikingly, the S6-HA linker straightened to rotate CaM away from the VSD, and the CaM-VSD interactions are replaced by S2-S3 linker interaction with PIP_2_ (15) (Fig. 1B). Functionally, KCNQ1 gating involves a two-step VSD activation, from the resting state to intermediate state, and finally to the activated state (8, 17–24). Each VSD activation step engages unique VSD-pore coupling interactions and triggers pore opening when the VSD adopts both the intermediate and activated states, leading to the IO and AO open states (8, 18, 21, 23, 24) (Fig. 1C). In this study, we provided functional and simulation analyses to connect the KCNQ1/CaM cryo-EM structures (14, 15) to the KCNQ1 voltage-dependent activation pathway.

Considering the activated VSD and the dilated pore in the CTD-straight conformation (15), the structure most likely reflects the AO state. Our MD network analysis revealed that VSD-pore coupling pathways specific to the AO state is emphasized in the CTD-straight structure (Fig. 3). Moreover, mutational disruption of CaM-VSD interactions, which is expected to promote the CTD-straight conformation, exhibited specific enhancement of channel transition into the AO state (Figs. 1, 2, 4). These results substantiate the suggestion that the CTD-straight conformation reflects to the AO state. While the CTD-straight structure was determined closer to physiological conditions with PIP_2_ (15), the CTD-bent structure was solved in the absence of PIP_2_ (14). This naturally raises the question of whether the CTD-bent structure exists in the physiological gating pathway. Our results show that mutational disruption the S2-S3 linker and CaM interface affects KCNQ1 gating and altered current (Figs. 1,2,4). Moreover, double mutant experiments found an interaction between the S2-S3 linker and CaM during gating (Fig. 2). These findings show that the CTD-bent conformation exists during voltage-dependent activation, and the CaM-VSD interface is functionally important for gating. Notably, the S2-S3 linker between the two cryo-EM structures interacts exclusively with either CaM or PIP_2_ (14, 15). Given that both structures exist during gating, our findings indicate that PIP_2_ and CaM compete for the S2-S3 linker during gating, such that the switch in the S2-S3 linker binding partner facilitates transition between the two conformations. Furthermore, our data suggest the exchange of CaM-VSD to PIP_2_-VSD interactions is related to VSD movement. Mutation of a PIP_2_-binding residue in the S2-S3 linker induced a depolarized shift in VSD movement (Supplementary Fig. 1B-C), demonstrating a direct relationship between VSD movement and PIP_2_ binding at the S2-S3 linker. The loss of the CaM-VSD interactions does not affect VSD movement (Fig. 4A-B), but selectively enhances AO state VSD-pore coupling and facilitates channel transition into the AO state (Figs 3F, 4C-E). Together these results suggest that VSD movement to the fully-activated state promotes PIP_2_ binding to the S2-S3 linker and the loss of CaM binding due to competition. CaM disengagement from the VSD subsequently triggers channel transition into the AO state. With this in mind, we propose that the CTD-bent structure reflects a transition state prior to PIP_2_ binding the S2-S3 linker, but the VSD has just moved to the fully-activated state. Although this transition state is likely transient during the gating process, the CTD-bent structure captured this state because it was solved at a depolarized voltage in the absence of PIP_2_.

In this study, we utilized MD simulations and network analysis to map a critical CTD hub which is allosterically downstream of the CaM-VSD interactions, and together the two interfaces affect channel transition into the AO state (Figs. 3 and 5-6). This CTD hub partially coincides with the S6-HA “RQKH” motif thought to be important for CTD rearrangement in the cryo-EM structural studies (15), but also contain HB-HC residues. We note that although the cryo-EM structures clearly show a large conformational change at the CaM-VSD interface, the identification of the CTD hub and its role in converting the channel to the CTD-straight conformation with an allosteric link to the CaM-VSD interactions are non-obvious from the structures. The MD network analysis approach thus proved to be a powerful tool to scrutinize structural data and guide functional experiments.

Based on the above conclusions, we propose a voltage-dependent KCNQ1 gating scheme integrating CaM and PIP_2_ (Fig. 6E) in which the VSD conformation controls CaM and PIP_2_ binding to the S2-S3 linker, providing conceptual improvement and structural basis to the transitions within the gating model. We emphasize that this gating model represents a starting point that accounts for all currently available structural and functional data. Although the model could quantitatively simulate the currents of KCNQ1 channels (8, 23, 24), more work is required to validate our proposed gating scheme.

When the VSD adopts the resting or the intermediate state, S2-S3 linker binds CaM, while PIP_2_ is present in the membrane (Fig. 6E: RC, IC, IO states). The CTD accordingly adopts the CTD-bent conformation. When the VSD transitions into the activated state, S2-S3 linker starts interacting with PIP_2_ which disengages CaM. The loss of CaM-VSD interactions then works together with the CTD hub (Fig 6E, green diamonds) to trigger transition into CTD-straight conformation and subsequent pore domain opening into the AO state.

We did not explicitly control for intracellular Ca^2+^ levels in this study. Several structural studies invariably revealed that the EF3 motif within CaM C-lobe is uncalcified over wide-ranging intracellular Ca^2+^ levels (14, 15, 33). Given that the CaM-VSD interactions mostly involved residues near EF3, we expect that CaM C-lobe adopted the same calcification state as seen in the structures with the oocytes intracellular Ca^2+^ levels in our experiments. How changing Ca^2+^ levels may affect the CaM-VSD interactions and KCNQ1 gating remains a subject for future studies.

PIP_2_ plays a vital role in our model by interaction with the S2-S3 linker to facilitate channel transition into the AO state. This is consistent with previous studies suggesting that PIP_2_ binds the S2-S3 linker (15, 36, 47–49). However, the S2-S3 linker may not be the only PIP_2_ binding site. As we showed in this study, PIP_2_ binding to the S2-S3 linker is VSD state-dependent (Supplementary Fig. 1B-C), indicating that PIP_2_ may bind elsewhere when the VSD is in the resting or intermediate states. Our previous studies found that PIP_2_ is required for KCNQ1 VSD-pore coupling in both the IO and AO states, which we attributed to PIP_2_ binding to the S4-S5 linker and the bottom of S5/S6 (36, 47).

These regions have also been suggested to bind PIP_2_ in other KCNQ channels (49). The S4-S5 linker, S5, and S6 may form a second PIP_2_ binding site specifically important for VSD-pore coupling in both the IO and AO states. This idea is further supported by our recent discovery of a PIP_2_-mimetic compound which enhances KCNQ1 VSD-pore coupling through occupancy of this second site (50). This second PIP_2_-binding site may be important for KCNQ1 VSD-pore coupling, while the S2-S3 linker PIP_2_ binding site is state-dependent and modulates channel transition into the AO state. Characterization of other PIP_2_ binding sites in KCNQ1 will be important in future studies.

With potential alternate PIP_2_ binding sites in mind, we note the caveat that the pore in the CTD-straight conformation, while dilated, is not fully open for current conduction (15). Thus, although the structure may capture the major features of the AO state, such as the loss of CaM-VSD interactions, the structure may reflect a conformation that is not exactly the AO state. The fact that the pore is not fully open could be because PIP_2_ in nanodiscs failed to occupy the second binding crucial for VSD-pore coupling. A second reason could be because the structure was solved without ATP, which is known to be required for KCNQ1 pore opening (51).

CaM has long been known to modulate voltage-gated ion channels by binding to their intracellular domains (6, 7, 26, 28, 31, 33, 52, 53). Yet, understanding how CaM modulates gating in context of the channel transmembrane domains has remained a challenge. Here, we show that the CaM-VSD interactions act as a switch to facilitate a CTD conformational rearrangement. This conformational change represents a striking difference, which may contribute to the unique IO and AO gating properties of KCNQ1. KCNQ1 is known to conduct predominantly at the AO state in the heart with KCNE1 association; but instead conducts primarily at the IO state in epithelial cells with KCNE3 association (8, 21). Modulation of KCNQ1 transition into the AO state thus represents a critical aspect of normal physiology. Here, we provide a mechanism for how CaM interacts with unexpected channel locations, such as the S2-S3 linker and the CTD hub, to control the AO state. Key structural interfaces contain residues linked to congenital LQTS mutations (13). The mutation R360G (S6), for example, is within the CTD hub and has been associated with LQTS. Thus, arrhythmias may result from dysfunction in this mechanism.

## Materials and Methods

### Site-directed mutagenesis

Point mutations were made in KCNQ1 channel and CaM utilizing overlap extension and high-fidelity PCR. DNA sequencing confirmed the presence of all mutants made. Mutant cRNA were made with the mMessage T7 polymerase kit (Applied Biosystems-Thermo Fisher Scientific).

### Channel expression in *Xenopus* oocytes

Stage V or VI oocytes were obtained from *Xenopus* laevis by laparotomy in accordance with the protocol approved by the Washington University Animal Studies Committee (Protocol #20190030). Oocytes were digested by collagenase (0.5-0.7 mg/ml, Sigma Aldrich, St Louis, MO) and injected with channel cRNAs (Drummond Nanoject, Broomall). Each oocyte was injected with cRNAs (9.2 ng) of WT or mutant KCNQ1. For experiments with KCNE1 or CaM co-expression, KCNE1 and CaM cRNAs were co-injected at 3:1 (KCNQ1:KCNE1) and 1:1 (KCNQ1:CaM) mass ratio, respectively. All VCF and some KCNQ1 constructs were injected with double the mass of cRNA to boost channel expression. Injected oocytes were incubated in ND96 solution (in mM): 96 NaCl, 2 KCl, 1.8 CaCl_2_, 1 MgCl_2_, 5 HEPES, 2.5 CH_3_COCO_2_Na, 1:100 Pen-Strep, pH 7.6) at 18°C for 2-6 days before recording.

### Two-electrode voltage clamp (TEVC) and voltage-clamp fluorometry (VCF)

Microelectrodes were made with thin wall borosilicate glass (B150-117-10, Sutter Instrument, Novato, CA) by a micropipette puller (P-97 or P-1000, Sutter Instrument, Novato, CA). The pipette resistance were 0.5-3 MΩ when filled with 3M KCl solution and submerged in ND96 solution. TEVC and VCF experiments were recorded in ND96 solutions at room temperature. Whole-cell currents were recorded with a CA-1B amplifier (Dagan, Minneapolis, MN) driven by Patchmaster (HEKA, Holliston, MA) software. Current recordings were sampled at 1 kHz and low pass filtered at 2 kHz. For VCF experiments, the KCNQ1 channels were labeled by incubating oocytes expressing KCNQ1 channels for 1 hour on ice in 10 μM Alexa 488 C5-maleimide (Molecular Probes, Eugene, OR) in high K^+^ solution in mM (98 KCl, 1.8 CaCl_2_, 5 HEPES, pH 7.6). The oocytes were washed and incubated in ND96 after labeling. VCF was performed by recording currents with the same instruments as TEVC. The fluorescence signals were measured by a Pin20A photodiode (OSI Optoelectronics, Hawthorne, CA) and an EPC10 patch clamp amplifier (HEKA, Holliston, MA) sampled at 1 kHz and filtered at 200 Hz. The EPC10 amplifier was controlled by the CA-1B amplifier to ensure simultaneous current and fluorescence measurements. All chemicals were obtained from Sigma Aldrich (St. Louis, MO).

### Electrophysiology data analysis

Data were analyzed with MATLAB (MathWorks, MA). The conductance-voltage (G-V) relationship calculation: the instantaneous tail currents following test pulses were normalized to the maximum current. The G-V relationship was fitted with a single Boltzmann equation in the form of

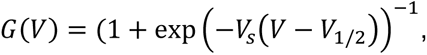

where *V* is the test pulse voltage, *V*_1/2_ is the half-activation voltage, and *V*_s_ controls the steepness of the Boltzmann equation. *V*_s_ is related to RT/zF, where *R* is the gas constant, *T* is the temperature, *z* is the equivalent valence, and *F* is the Faraday constant. The fluorescence-voltage (F-V) relationship calculation: linear photobleaching correction was applied to each fluorescence trace by subtracting the line obtained by linear fitting of the 2-seconds fluorescence signal prior to the start of the test pulse. The F-V relationship was calculated using fluorescence signals at the end of the test pulses and normalized to the maximum fluorescence signal change. The F-V relationship was then fitted with a double Boltzmann distribution in the form of

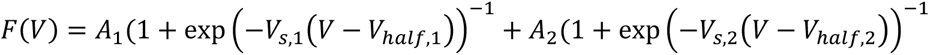

where *V*_half,1_ and *V*_half,2_ correspond to the half-activation voltages for *F*_1_ and *F*_2_, respectively. Other variables are similar defined as in the G-V relationship fits. One-way analysis of variance (ANOVA) followed by Tukey’s HSD test was utilized to compare *V*_1/2_ values from multiple mutants (Figs 1C, 4B). All other statistical significance calculations were performed with Student’s T-test.

Current activation kinetics were quantified by fitting the raw 4-seconds test-pulse currents to a bi-exponential function in the form of

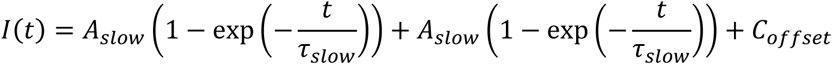

Each current trace was baseline corrected using the mean current at the holding potential for each trace. The fast current component was first estimated by fitting the first 0.5 seconds of the current. A second overall fit was then applied to the entire 4-seconds test pulse current, with the fast time constant constrained within ±25% of the first fit. Each trace was then normalized by the fitted steady-state amplitude (*A*_fast_ + *A*_slow_ + *C*_offset_) to allow comparison between different cells. The fraction of total current carried by the fast component was calculated with the equation *A*_fast_ / (*A*_fast_ + *A*_slow_).

### Molecular dynamics simulation

The human CTD-bent structure (PDB ID 6UZZ) and the CTD-straight without KCNE3 (PDB ID 6V01) (15) were simulated in equilibrium with three replicas amounting up to ∼1*μ*s per system. Similar to the 1*μ*s trajectory from (54), we used a POPC membrane, as well as explicit solvent neutralized with 0.1 M KCl. Missing loops were modeled with modeller and the DOPE score (55). The equilibration was done according to the standard 6-step Charmm-gui procedure (56), extending the sixth step to 3 ns.

The three simulations of the *Xenopus* structure (14), were launched from the minimized and equilibrated system, i.e. the first frame, of the already published 1*μ*s production trajectory (54), by generating new velocities according to a Maxwell distribution. Each new simulation was equilibrated for 375 ps while reducing restraints on the protein atoms, according to the standard 6-step Charmm-gui procedure (56).

Each system was simulated in an NPT ensemble with a 2 fs time step. Temperature was held constant at 303.15 K by the Nosé-Hoover thermostat (57) during production and the V-rescale thermostat (58) during equilibration. Pressure was maintained at 1 bar with the semi-isotropic Parrinello-Rahman barostat (59) during production and the Berendsen barostat (60) during equilibration. Short-ranged electrostatics were computed with a 1.2 nm cutoff, such that the switching function started at 1.0 nm, while long-ranged electrostatics were modeled with PME (61). Moreover, Charmm36 force field (62) and Gromacs 2018 (63) were used, water was modeled with TIP3 and bonds involving hydrogens were constrained with LINCS (64). The total simulation time of the three trajectories was 1.2 *μ*s. The simulation parameters of the 1 *μ*s long trajectory of the xenopus structure (14) are previously described in (54). This was simulated in equilibrium with Gromacs 2016 (63).

The RMSD along each trajectory was calculated with MDtraj 1.9.3 (65). Structural visualization was done with VMD (66).

### Creating a residue-residue network

#### Semi-binary contact map

Because continuous contact maps yield more robust networks than binary (38), we used a continuous contact map, such that distances below *c* = 0.45 nm were weighted by 1, and larger distances were down-weighted continuously until a cutoff, *d_cut_* = 0.8 nm, where the interactions were deemed insignificant. This was done with a truncated Gaussian kernel which is *K*(*d)* =1 if *d* ≤ *c* and 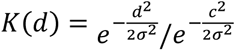. if *d* > *c* The denominator, 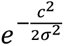, makes sure that the contact-kernel is continuous at *d* = *c*. To determine *σ*, we used *K* (*d_cut_* = 0.8) = 10^-5^so that 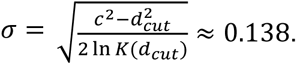

The semi-binary contact between residues *s_i_* and *s_j_* from all frames were averaged to form the final semi-binary contact map,

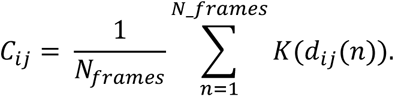

#### Mutual information

Mutual information (MI) was used to estimate correlation of residue movements. The MI, *I*, between residue *s_i_* and *s_j_* was estimated based on distances from their respective equilibrium positions (67). The residue position was taken as the centroid of the residue, thus accounting for both side chain and backbone motions. MI is calculated as

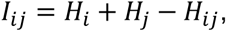

where *H_i_*is the entropy of residue *s*_*i*_,

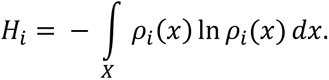

The density was estimated with Gaussian mixture models (GMM) (68–70) with the GMM implementation in Scikit-learn 0.21.2 (71), and the framework presented in (68, 69). The number of Gaussian components was chosen with the Bayesian information criterion (72), varying between 1 and 5 components.

The number of frames not only affects the density estimate, but also the approximated entropy. To increase accuracy, parametric bootstrapping was done by repeatedly drawing *N_frames_* new samples from the estimated density. Using the new samples, new MI matrices were estimated. This was repeated 10 times. The final MI matrix was taken as the average of all estimated matrices.

### Current flow analysis

The MI matrix and semi-binary contact map make up the full residue network with adjacency matrix elements *A_ij_* =*Cij I_ij_*

#### Definition of current flow betweenness

Assuming that information spreads from source residues *S*_0_ to sink residues *S*_1_ on the network according to diffusion, or equally a random walk, where the sink nodes are absorbing, we can use current flow betweenness (referred to as information flow in this paper) to describe the allosteric pathways.

First, the network Laplacian is computed as, *L* = *D*-*A*, where *D_ij_* = ∑_j_ *A_ij_* is the diagonal degree matrix. The inverse reduced Laplacian, 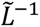, is obtained by removing absorbing nodes (i.e. sink nodes) from the Laplacian, inverting this matrix, and then again including zero-rows and columns at the absorbing nodes.

Given a supply vector, 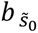 which represented an injection of one unit information (or current) in a single source node 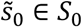 that would exit in the sink nodes, the information flow through an inner node (not source or sink), or residue, *s_i_* was calculated as

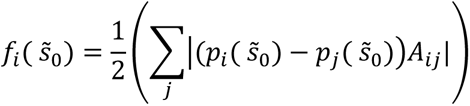

Where 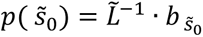. The information flow through the source and sink nodes were set to zero to only consider inner states in the analysis. The final information flow was averaged over the source nodes of one subunit,

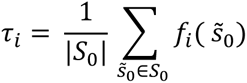

#### Symmetrization over subunits

The current flow through each residue was calculated for sources in each subunit, one at a time. The resulting current flow profiles were centered on the first subunit. The current flow profiles were then replicated over the structure, and the total current flow through one residue was taken as the average contribution from each subunit. The average and standard deviation over the structure were calculated from the replicated current flows.

#### Identifying important residues via hubs

To reveal the communication between the VSD and the pore and highlight hubs with important residues, two different pairs of sources and sinks were studied:

1. Gating charges on S4 (R1, R2, Q3, R4)-S349
2. CCaM (residues 82-148 in CaM, the C-term domain)-S349

This analysis was done on the CTD-bent structure with source/sink pairs 2 vs. 1, and on CTD-straight vs. CTB-bent using source/sink pairs 1. To highlight differences, we compared the information flow profiles with simple subtraction of information flow of mutual residues (ie. residues that are present in both structures). The standard deviation of the delta-information-flow was calculated with the square root of the added variance of each information flow profile.

#### Correlation of information flow profiles

The results from the Xenopus KCNQ1 simulations were compared to human KCNQ1 simulations by first converting the Xenopus residue numbering to the human structure. Linear regression was performed and the Pearson correlation coefficient computed. This was done using the information flow of mutual residues. The linear regression was done with Scikit-learn 0.21.2 (71).

### Residue interaction analysis

Interactions were quantified by the minimum distance between all heavy atoms of the two residues along the second half of each trajectory. The interaction distances were calculated using the python package MDtraj 1.9.3 (65). The distance densities over the trajectory were estimated with GMM, with the number of components chosen by Bayesian information criterion by varying between 1 and 6 components. The average density was scaled between 0 and 1. The standard error was calculated over the four subunits.

## Acknowledgments

This work was supported by grants from NIH R01HL126774 (J.C.), R01NS 092570 (J.R.S.), grants from the Science for Life Laboratory and the Göran Gustafsson Foundation (L.D.). The simulations were performed on resources provided by the Swedish National Infrastructure for Computing (SNIC) at the PDC Centre for High Performance Computing (PDC-HPC).

## Author Contributions

P.W.K and J.C. initiated this work. J.C. and L.D. directed the approaches used. The experimental work was conducted by P.W.K., J.S., K.M.W., A.K.D., and A.H.C. The computational work was conducted by A.M.W. All authors participated in data analysis. P.W.K., A.M.W. J.C. and L.D. wrote the paper with input from all authors.

